# OpenCafeMol with 3SPN.2 DNA model: GPU Acceleration for Long-Time Coarse-Grained Chromatin Simulations

**DOI:** 10.64898/2026.03.18.712524

**Authors:** Masataka Yamauchi, Yutaka Murata, Toru Niina, Shoji Takada

**Author notes:** Contributed equally (Masataka Yamauchi, Yutaka Murata).

## Abstract

There is a growing demand for molecular dynamics simulations to explore longer timescale behavior of giant protein-DNA complexes such as chromatin. To address this need, we extended OpenCafeMol, a GPU-accelerated residue-level coarse-grained molecular dynamics simulator originally developed for proteins and lipids, to support 3SPN.2 and 3SPN.2C DNA models. We also implemented a hydrogen-bond-type many-body potential to model DNA-protein interactions more accurately. To further improve computational efficiency, we introduced a localized scheme for calculating base-pairing and cross-stacking interactions. Benchmark tests show that OpenCafeMol on a single GPU achieves up to 200-fold speed-up for DNA-only systems and up to 100-fold speed-up for DNA-protein complexes compared to CPU-based simulations. To demonstrate the capability of our implementation for long-timescale biological processes, we simulated an archaeal SMC-ScpA complex undergoing DNA translocation via segment capture (a proposed mechanism for DNA loop extrusion) in the presence of a DNA-bound obstacle. We observed continuous captured-loop growth accompanied by obstacle bypass within the segment capture framework.

## INTRODUCTION

Coarse-grained (CG) molecular dynamics (MD) simulations have become an indispensable tool for investigating long-time dynamics of complex biomolecular systems. By simplifying molecular representations while preserving key physicochemical features^1,2^, CG models enable simulation of processes that are computationally prohibitive for all-atom models. Over the past decades, CG approaches have substantially advanced our understanding of protein folding, aggregation, phase separation, molecular machinery, and membrane-related phenomena^3,4^.

In parallel, advancements in high-performance computing, particularly the use of graphics processing units (GPUs), have significantly transformed MD simulations. Widely used MD software packages, including GENESIS^5–7^, GROMACS^8^, Amber^9–11^, OpenMM^12^, NAMD^13^, and LAMMPS^14^ now support GPU acceleration. These performance gains extend accessible timescales and enable more comprehensive sampling of the conformational landscapes and kinetics of biomolecular systems.

Leveraging GPU acceleration for CG simulations provides a practical route to meet a growing demand for simulating larger and more intricate biomolecular systems over biologically relevant timescales. Recently, our group developed OpenCafeMol, a MD simulator designed for residue-resolution CG models of proteins and lipids^15^. Built on the OpenMM framework, OpenCafeMol achieves efficient performance across various hardware platforms, including GPUs. Benchmark tests have demonstrated that, on a single high-end GPU, protein and lipid membrane simulations are accelerated by up to 100-fold and 240-fold, respectively, compared to CPU-based simulations. However, while OpenCafeMol currently supports the AICG2+ CG protein model^16^, the hydrophobicity scale (HPS) model^17,18^, and the iSoLF lipid model^19^, it has lacked support for nucleic acids. Integrating a CG DNA model into OpenCafeMol is thus a crucial step to broaden its applicability to biologically significant processes, particularly those involving chromatin dynamics.

In this study, we extended OpenCafeMol by integrating the 3SPN.2 and 3SPN.2C CG DNA models^20,21^ using the OpenMM library. The 3SPN (three-site-per-nucleotide) model^22^, originally developed by the de Pablo group, represents each nucleotide with three particles: one for the phosphate group, one for the sugar, and one for the nucleobase, reproducing DNA’s structural and thermodynamic properties. The 3SPN.2^20^ model reproduces major structural and thermodynamic properties of DNA, while the refined 3SPN.2C^21^ incorporates sequence-dependent bending propensity, thereby providing an even more faithful representation of DNA conformational landscapes. In addition to incorporating the 3SPN.2 and 3SPN.2C models, we implemented a hydrogen-bond-type potential^23^ to more accurately represent orientation-dependent protein-DNA interactions. These new features are fully compatible with the AICG2+ CG protein model. Indeed, the combined use of 3SPN.2 or 3SPN.2C with AICG2+ and hydrogen-bond interactions has been successfully applied to a variety of DNA-protein complexes^24^, including nucleosomes^25–27^, RNA polymerase II^28^, DNA replication machinery^29,30^, and structural maintenance of chromosomes (SMC) complexes^31^.

GPU acceleration of 3SPN.2 DNA models using OpenMM poses specific challenges, especially due to their intricate base-pairing and cross-stacking interactions. In a previous work by Open3SPN2 ^32^, these many-body interactions implemented via CustomHBondForce in OpenMM were computationally expensive, yielding limited performance gains on GPUs. Here, we demonstrate that optimizations in CustomHBondForce introduced in OpenMM 8.1^33,34^ alleviate this bottleneck. Furthermore, to maximize efficiency, we introduce a localized interaction scheme that restricts base-pairing and cross-stacking calculations to relevant neighbors. These approaches drastically reduce computational costs, achieving the substantial speed-ups required for long-timescale chromatin simulations

## METHODS

### 3SPN.2 DNA model

The 3SPN.2 and 3SPN.2C models represent each nucleotide using three particles corresponding to the phosphate, sugar, and base groups. Each particle is positioned at the center of mass of its respective molecular moiety (Fig. 1). This simplified representation reproduces the structural and thermodynamic properties of DNA at a reasonable computational cost. The potential energy, 𝑉_3SPN.2_, is expressed as a sum of eight individual terms:

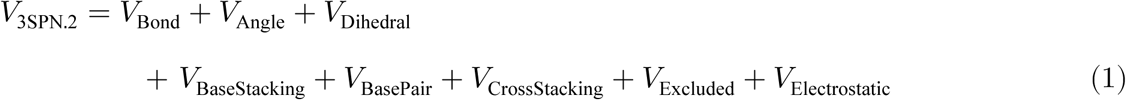

**Figure 1.**
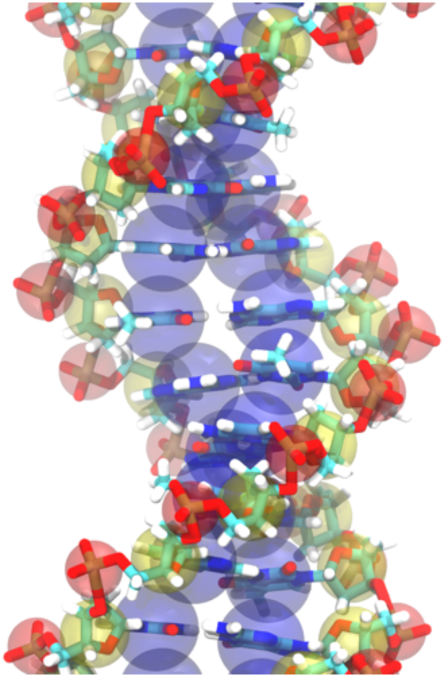
Mapping particles from all-atom to coarse-grained representation of nucleic acids. Each nucleotide is represented with three particles: phosphate (red), sugar (yellow), and base (blue). Each particle is placed at the center of mass of its respective molecular moiety.

The first three terms describe bonded interactions, including bond stretching, angle bending, and torsion angles. The remaining five terms account for non-bonded interactions. Among these, 𝑉_Stacking_, 𝑉_BasePair_, and 𝑉_CrossStacking_ terms represent base-base interactions, while 𝑉_Excluded_ and 𝑉_Electrostatic_ represent pairwise interactions.

### Bonded terms

The bond stretching term describes the interaction between two adjacent connected sites (e.g., phosphate-sugar or sugar-base), which is expressed as a sum of quadratic and quartic terms as follows:

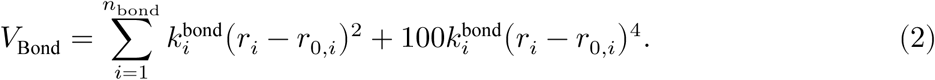

Here, 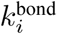 and 𝑟_0_,*_i_* denote the force constant and equilibrium bond length, respectively. The equilibrium distances are set to fixed values in the 3SPN.2 model^20^, whereas in the 3SPN.2C model, they are computed from a fiber structure of B-form DNA generated using X3DNA^35,36^.

The angle bending term is represented by a harmonic potential as a function of the angle 𝜃_𝑖_ formed by three consecutive sites.

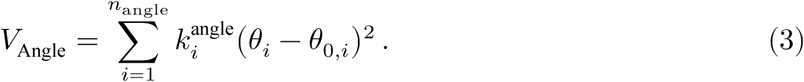

Here, 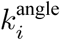 and 𝜃_0_,*_i_* denote the force constant and equilibrium angle, respectively. The equilibrium angles are computed from a fiber structure of B-form DNA, which can be generated using X3DNA^35,36^.

The dihedral term consists of a Gaussian term and a cosine potential, which is given by:

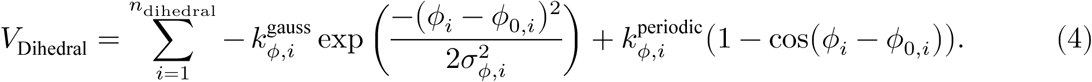

Here, 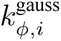 is the well-depth of Gaussian potential, 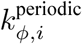 is the height of the periodic cosine potential, 𝜙_0,𝑖_ is the equilibrium dihedral angle, and 𝜎_𝜙,𝑖_ is the well-width of Gaussian potential. The 3SPN.2 model uses only the Gaussian potential^20^ while the 3SPN.2C model uses a combination of Gaussian and cosine potentials^21^. The Gaussian potential is applied for the dihedral angles between sugar and phosphate particles, whereas the cosine potential is applied for both sugar-phosphate dihedrals and those involving base particles to introduce sequence-dependent torsional effects. For the 3SPN.2 model, equilibrium angles are computed from a fiber structure of B-form DNA, whereas in the 3SPN.2C model they are sequence-dependent, derived from a template based on geometric parameters. Reference structures for both 3SPN.2 and 3SPN.2C can be generated using X3DNA^35,36^.

### Base-base non-bonded terms

The non-bonded interactions between nucleobases consist of base-stacking, base-pairing, and cross-stacking, which act depending on the distances between the nucleobases as well as specific orientation-dependent angles. The distance dependence is modeled with the Morse potential, which is divided into repulsive and attractive components as follows:

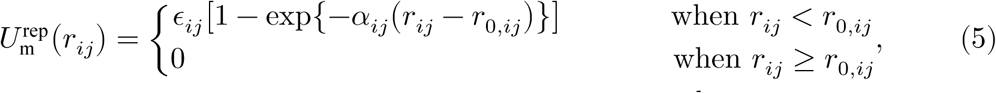

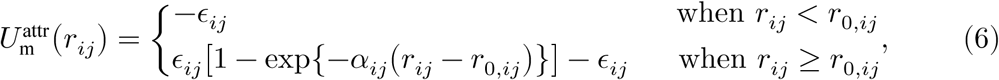

where 𝜖_𝑖𝑗_ is the depth of the well of the attraction between particles 𝑖 and 𝑗, 𝑟_0,𝑖𝑗_ is the equilibrium separation distance, and 𝛼_𝑖𝑗_ is a parameter that controls the range of attraction.

To represent the specificity of the interactions, orientation dependence is incorporated into the attractive component via a filter function defined as:

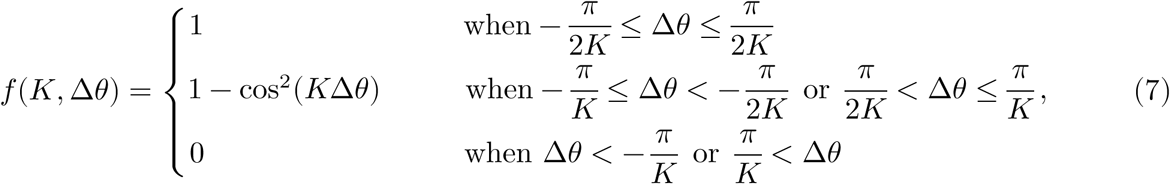

where Δ𝜃 = 𝜃 − 𝜃_0_ represents the deviation in the angle that represents the orientation. The 𝜃_0_ is the corresponding angle at the ideal B-form DNA. The parameter 𝐾 defines the width of the allowed angle range.

The base-base stacking energy, which is the interaction between the two consecutive bases of the same strand, is represented as follows:

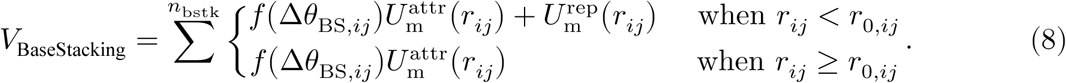

Here, 𝑟_𝑖𝑗_ is the distance between two adjacent base particles of the same strand (Fig. 2A). 𝜃_BS_ is the angle between a vector connecting a sugar and base site within the same nucleotide and the vector connecting the current base to its neighbor base (Fig. 2A), which represents the orientation dependence of the base stacking.

**Figure 2.**
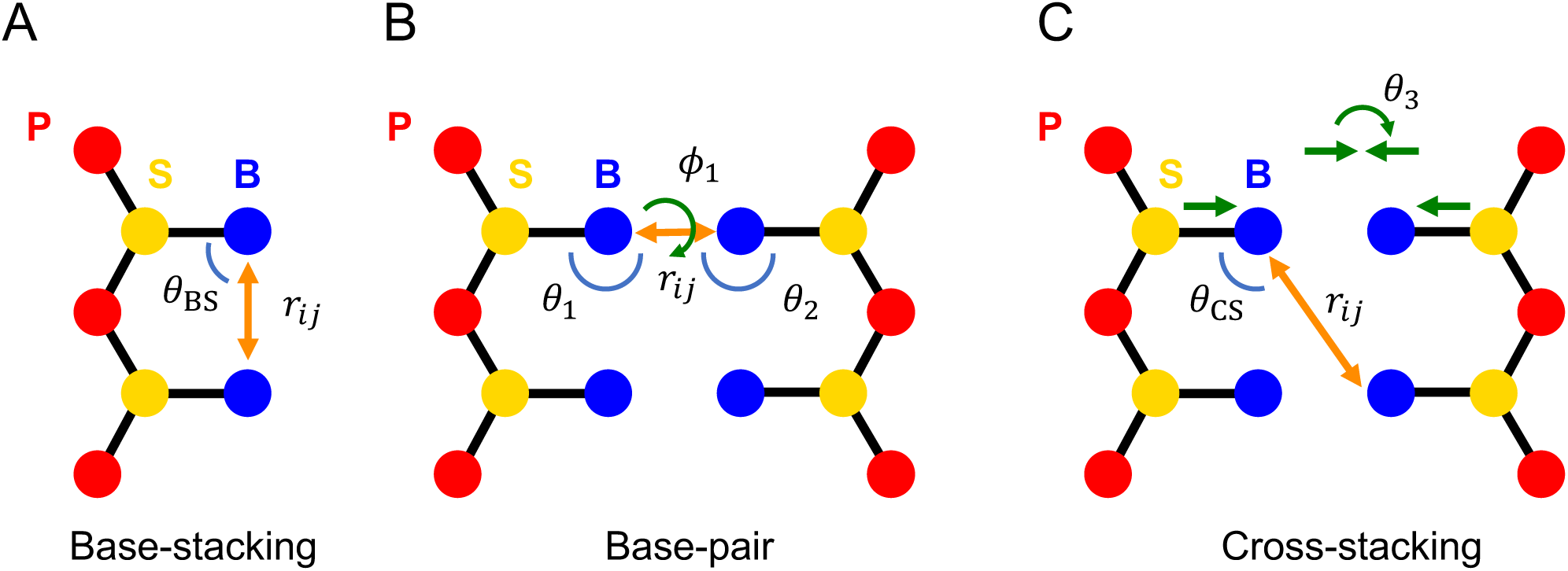
Definition of variables used for (A) intra-strand base-stacking term, (B) base-pairing term, and (C) cross-stacking term.

The base-pair energy represents the interactions between base pairs of the double strands connecting through hydrogen bonds, which is represented as follows:

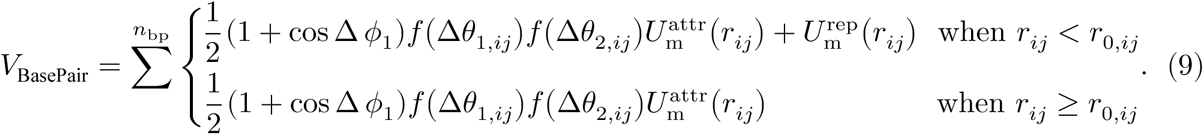

Here, 𝑟_𝑖𝑗_ is the distance between two bases connecting through the hydrogen bonds (Fig. 2B). The orientation dependence is introduced through two angles, 𝜃_1_ and 𝜃_2_, and one dihedral angle 𝜙_1_ (Fig. 2B). The angle 𝜃_1_ is defined as the angle between a vector connecting the sugar and the base of 𝑖-th nucleotide and the vector connecting the same base of 𝑖-th nucleotide and its hydrogen-bonding base of 𝑗-th nucleotide. The angle 𝜃_2_ is defined as the angle between a vector connecting the sugar and the base of 𝑗-th nucleotide and the vector connecting the same base of 𝑗-th nucleotide and its hydrogen-bonding base of 𝑖-th nucleotide. The dihedral angle 𝜙_1_ is defined by the four particles: the sugar and the base of 𝑖-th nucleotide and the base and the sugar of 𝑗-th nucleotide.

The cross-stacking energy represents the base-base interaction between the base particle of the 𝑖-th nucleotide in one strand and the base particle of the 𝑗-th nucleotide which is adjacent to the base particle forming a hydrogen bond with the 𝑖-th nucleotide (Fig. 2C). Because the distance between 𝑖-th and 𝑗-th bases, 𝑟_𝑖𝑗_, cannot be close, only the attractive part of the Morse potential is included as follows:

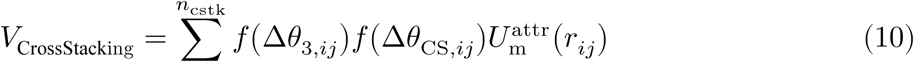

where the orientation dependence is introduced through two angles, 𝜃_3_ and 𝜃_CS_ (Fig. 2C). The angle 𝜃_3_ is the angle between the vector connecting the sugar and the base sites of 𝑖-th nucleotide and the corresponding vector of the nucleotide that makes a hydrogen bond with 𝑖-th nucleotide (note that this is not the 𝑗-th nucleotide). The angle 𝜃_CS_ is the angle between the vector connecting the sugar and the base sites of 𝑖-th nucleotide and the vector connecting the 𝑖-th and 𝑗-th base sites which are in the cross-stacking relation. In the original literature of 3SPN.2 model^20^, the cross-stacking is defined asymmetrically, requiring an *a priori* assignment of “sense” and “antisense” strands. This definition introduces a physical inconsistency, as the calculated potential energy varies depending on the arbitrary strand labeling. Furthermore, such strand dependence becomes problematic in simulations involving strand hybridization or nick formation, where the sense/antisense distinction may become ambiguous or dynamically change during the simulation. To address this, OpenCafeMol^15^ (similar to Mjolnir^37^) adopts a symmetrized cross-stacking potential, ensuring that the energy remains invariant under strand exchange.

### Pairwise non-bonded terms

The excluded volume interactions among phosphate, sugar, and base particles contain the repulsive section of a Lennard-Jones potential, which is as follows:

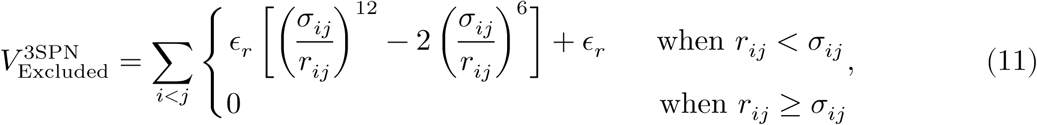

where, 𝜖_𝑟_ is the energy parameter, 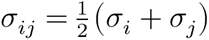 is the average particle diameter, and 𝑟_𝑖𝑗_ is the distance between particles 𝑖 and 𝑗. The excluded volume interactions are not applied between particles involved in bonded interactions (𝑉_Bond_, 𝑉_Angle_, and 𝑉_Dihedral_), base-pairing interactions (𝑉_BasePair_), or particles belonging to adjacent nucleotides within the same strand.

The electrostatic interaction is calculated under Debye-Hückel approximation, which can be written as follows:

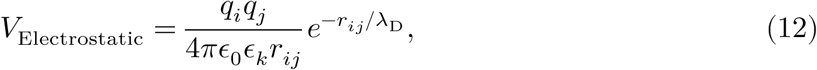

where, 𝑞_𝑖_ and 𝑞_𝑗_ are charges of particles 𝑖 and 𝑗, 𝜖_0_ is electric constant, 𝜖_𝑘_ is dimensionless dielectric constant, 𝑟_𝑖𝑗_ is the distance between two non-bonded charged particles. 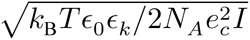 is Debye length, where 𝑘_𝐵_ is Boltzmann constant, 𝑁_𝐴_ is Avogadro’s number, 𝑒_𝑐_ is the elementary charge, and 𝐼 is the ionic strength of the solution. The 3SPN.2 and 3SPN.2C models use -0.6 charge at the phosphate group taking into consideration of Manning-like counter ion condensation.

### DNA-protein interactions

#### Protein-DNA excluded volume

To prevent overlap between protein and DNA particles, excluded volume interactions are applied using the same formulation as in the AICG2+ CG model^16^:

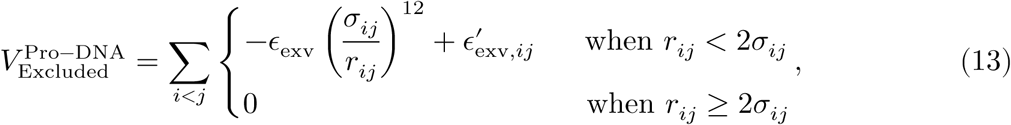

where 𝜎_𝑖𝑗_ represents the excluded volume distance, which depends on particle type. The parameter 𝜖_exv_ is force coefficient and set to 0.6 kcal/mol, and 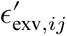 is a correction for the cutoff which corresponds to (1/2)^12^𝜖_exv_.

#### Protein-DNA electrostatic interaction

Electrostatic interactions between protein and DNA particles are calculated under the Debye-Hückel approximation, as described in Eq. (12). While the partial charge of phosphate particles is set to -0.6 in DNA-DNA interactions, this charge may not be optimal for DNA-protein interactions. Based on our previous investigation^38^, we recommend assigning a charge of -1.0 to the phosphate group for DNA-protein interactions.

#### DNA-protein hydrogen bond-type interaction

Hydrogen bond-type interactions between basic amino acid residues and the phosphate groups of the DNA backbone are essential for stabilizing DNA-protein complex structures. For example, a previous CG simulation study successfully reproduced the sliding dynamics of nucleosomes by incorporating both long-range electrostatic interactions and short-range hydrogen bond interactions between the histone octamer and DNA, while simulations incorporating only electrostatic interactions did not^23^.

In CG modeling, hydrogen bonds can be represented using a potential function that accounts for distance and angle dependencies, as described below:

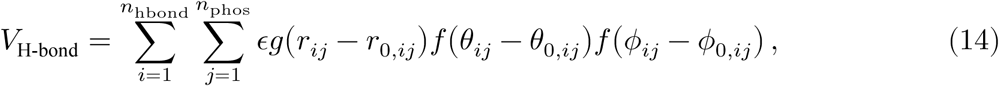

where the first summation runs over the list of hydrogen bonds identified from reference crystal structures or all-atom simulations, and the second summation runs over all DNA phosphate particles. 𝜖 is the energy parameter that determines the strength of the hydrogen bond. 𝑟_𝑖𝑗_ is the distance between a C_α_ particle of 𝑖-th hydrogen-bond-forming residue and 𝑗-th phosphate particle (Fig. 3). 𝜃_𝑖𝑗_ is the angle between the vector connecting the C_α_ particle of 𝑖-th the hydrogen-bond-forming residue to the phosphate and the vector connecting the two C_α_ particles neighboring the hydrogen-bond-forming one along the polypeptide chain (Fig. 3). 𝜙_𝑖𝑗_ is the angle between the hydrogen-bond-forming residue, the hydrogen-bond-forming phosphate, and the sugar particles that is in the same nucleotide of the phosphate (Fig. 3). The parameters with the subscript 0, namely 𝑟_0,𝑖𝑗_, 𝜃_0,𝑖𝑗_, and 𝜙_0,𝑖𝑗_, represent the representative values of distance and angles for 𝑖-th hydrogen bond, which are derived from reference crystal structures or all-atom simulations. The angular dependence is controlled by a filter function 𝑓, as defined in Eq. (7) while the distance dependence is modeled using a Gaussian function which is defined as:

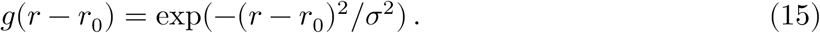

**Figure 3.**
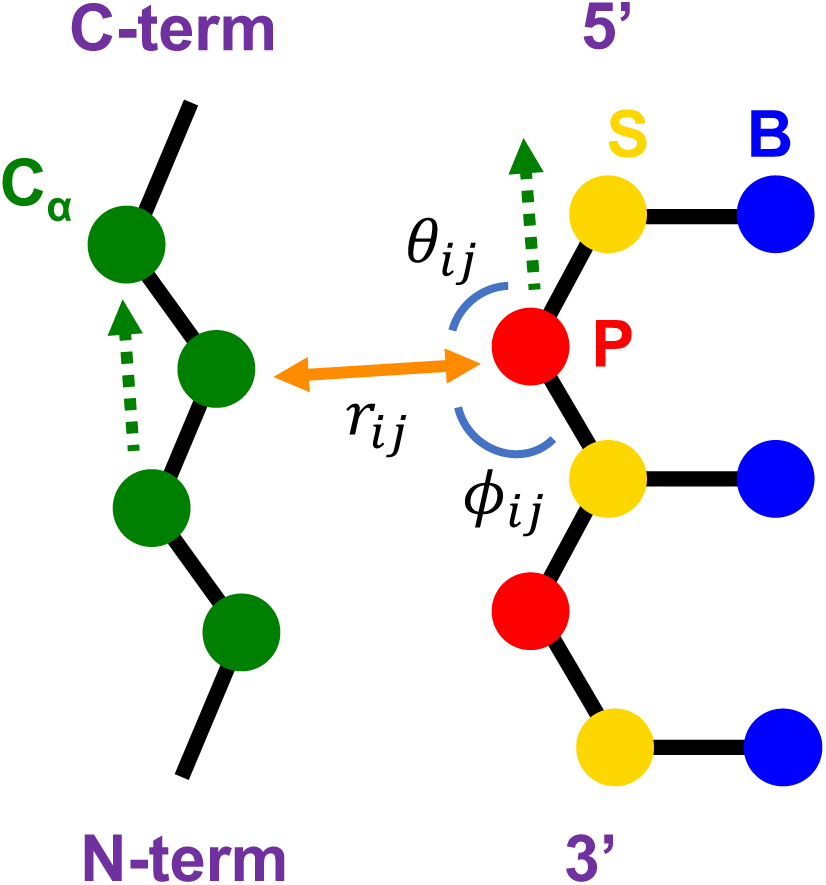
Definition of the variables used for coarse-grained hydrogen-bond between protein and DNA.

The potential widths 𝜎, Δ𝜃, and Δ𝜙 are typically set to 1 Å and 10 degrees, respectively, ensuring specificity in hydrogen bond distances and angles.

### Implementation

OpenCafeMol is a simulation package designed for efficient residue-resolution CG MD simulations, which builds upon the OpenMM C++ API^15^. Within OpenCafeMol, each potential energy term is managed by a dedicated “ForceFieldGenerator” (FFG) class, which internally utilizes OpenMM’s force classes. This object-oriented design of the software enables the users to easily incorporate new force fields.

The 3SPN.2 model, 3SPN.2C model, and the hydrogen-bond-type interactions involve more intricate potential function forms than the standard potentials commonly used in biomolecular simulations. To accommodate this complexity, we utilized OpenMM’s "custom" force classes that allow developers to define arbitrary algebraic expressions directly as strings. OpenMM’s just-in-time compiler then generates highly optimized kernels for various hardware backends, including CUDA and OpenCL, thereby relieving users from the burden of low-level optimization^39,40^. Table 1 lists all the implemented OpenCafeMol FFG classes and their corresponding OpenMM classes.

**Table 1.**
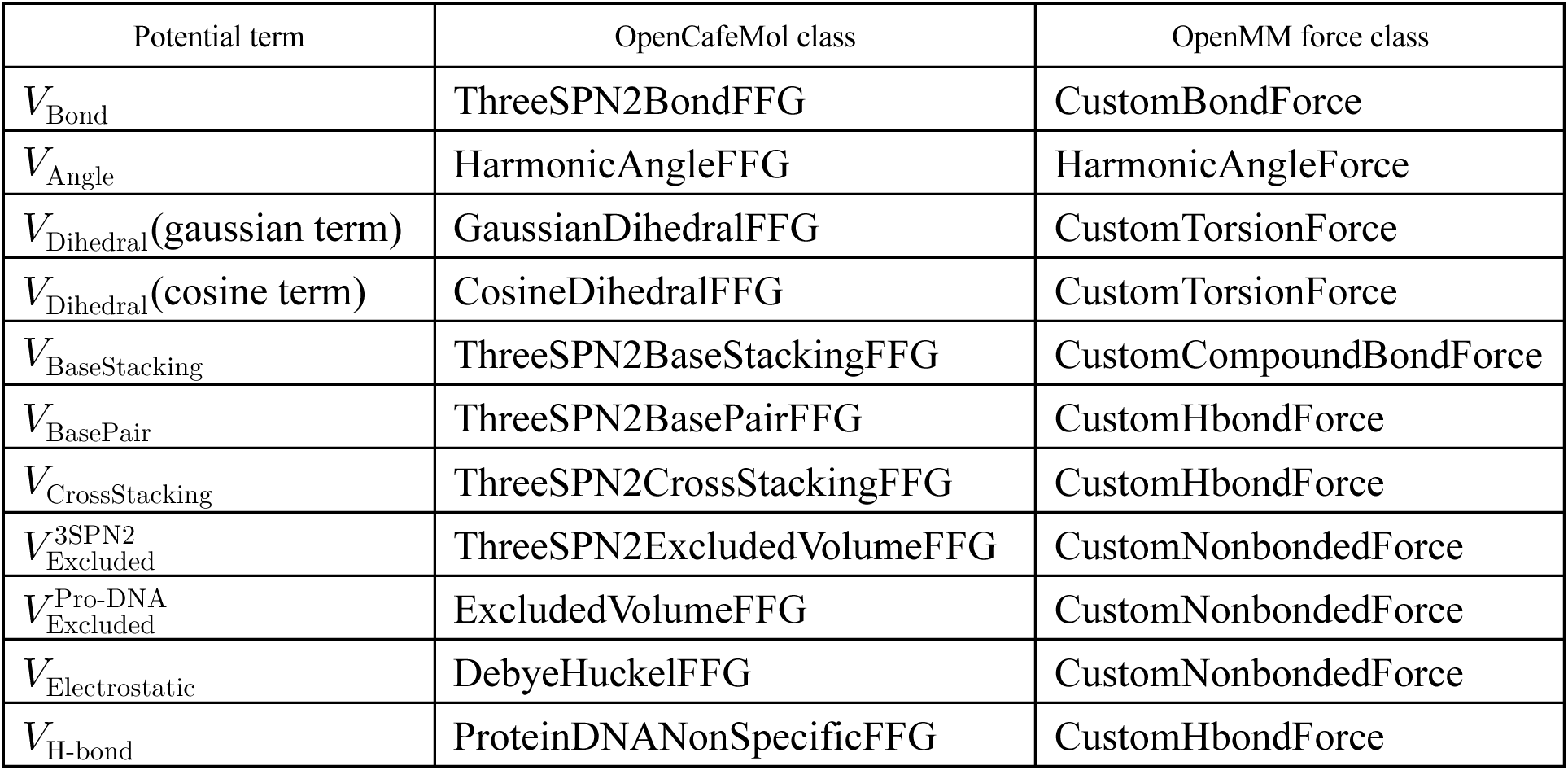
OpenCafeMol classes for the 3SPN2 and 3SPN.2C DNA model.

In addition to implementing the FFG classes for the 3SPN.2, 3SPN.2C, and hydrogen-bond-type interactions, we also introduced specialized FFG classes to address the high computational cost of many-body interactions. The standard implementation of the base-pairing (𝑉_BasePair_) and cross-stacking (𝑉_CrossStacking_) interaction relied on CustomHbondForce. This approach inherently involves iterating over all possible donor-acceptor pairs, leading to an 𝑂(𝑁^2^) scaling with respect to the number of nucleotides^34^. However, for simulations maintaining a double-strand DNA topology, these interactions are physically significant only between Watson-Crick base pairs and their immediate neighbors. This means that enforcing these interactions across all nucleotide pairs is unnecessary and wasteful as long as the DNA remains double-stranded. Therefore, there is room to improve computational efficiency by restricting the calculation to pairs that are known *a priori* to form base pairs and their adjacent nucleotides.

To eliminate this bottleneck, we implemented a localized interaction scheme using CustomCompoundBondForce (Table 2). This method defines 𝑉_BasePair_ and 𝑉_CrossStacking_ strictly between predetermined nucleotide pairs. Consequently, to use these functions, users must specify in the input file which nucleotide pairs will engage in base-pairing and cross-stacking interactions. This approach improves performance by eliminating the redundant calculation of all possible pairs. Reflecting this localized treatment of the interactions, we have named these new FFGs as ThreeSPN2BasePairLocalFFG and ThreeSPN2CrossStackingLocalFFG. Note that this localized scheme assumes the conservation of base-pairing and cross-stacking (i.e., double-stranded DNA topology). Therefore, it is primarily intended for simulations where the overall DNA topology remains fixed during the simulations. Although minor local fluctuations, such as transient bubble formation, could be handled by registering potential adjacent interacting pairs *a priori*, the standard global interaction scheme must be employed to capture spontaneously forming interactions in processes involving dynamic structural changes, such as hybridization, strand exchange, or defect formation where non-canonical base-pairing may occur.

**Table 2.**
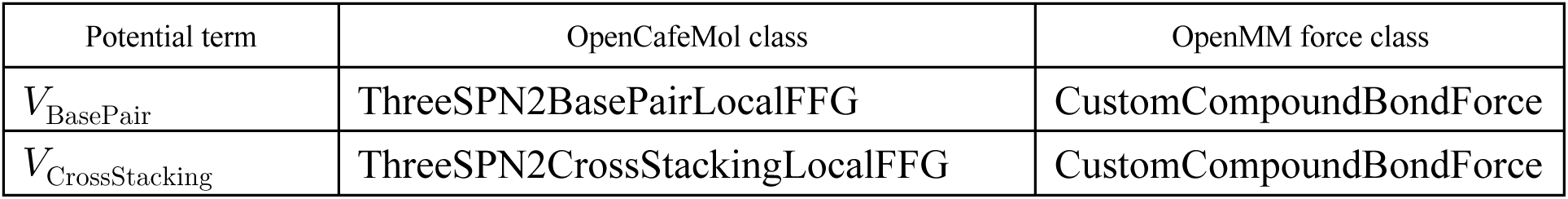
OpenCafeMol classes for localizing base-pair and cross-stacking interactions.

| Potential term | OpenCafeMol class | OpenMM force class |
| --- | --- | --- |
| $V_{\text{BasePair}}$ | ThreeSPN2BasePairLocalFFG | CustomCompoundBondForce |
| $V_{\text{CrossStacking}}$ | ThreeSPN2CrossStackingLocalFFG | CustomCompoundBondForce |

## COMPUTATIONAL DETAILS

All CPU-based simulations were performed using CafeMol version 3.2.1^41^ (https://www.cafemol.org). Langevin dynamics was adopted for time integration^42^ of the MD with a friction coefficient set to the default value of CafeMol. The time step was set to 0.2 café-time, where a time unit of 1 café-time corresponds to 49 fs. All CPU-based benchmarks were carried out with Intel Xeon CPU MAX 9490 on the supercomputer system of Academic Center for Computing and Media Studies at Kyoto University.

All GPU-based simulations were performed using OpenCafeMol^15^ which builds on the OpenMM library. We examined different OpenMM versions (7.7.0 and 8.1.1)^33,39^ to investigate how optimizations introduced in a recent major update of OpenMM8 affect performance. Langevin integrator implemented in OpenMM was used for time integration. The friction coefficient was set to 0.01 café-time^-1^. The time step was set to 0.2 café-time. All GPU-based benchmarks were carried out with NVIDIA RTX4090 graphics card.

## RESULTS AND DISCUSSION

### Validation of Potential Energy Values

To validate the implementation of the 3SPN.2 and 3SPN.2C CG DNA models in OpenCafeMol, we compared the energy values for each potential term calculated by OpenCafeMol with those obtained with previously developed software, CafeMol^41^ and Mjolnir^37^. A double-stranded DNA molecule with the sequence ATACAAAGGT GCCGAGGTTTC TATGCTCCCACG was used for this validation. 100 conformations were generated through short MD simulations using CafeMol. Figures 4 and 5 show the comparison of each potential energy term between OpenCafeMol and CafeMol for the 3SPN.2 and 3SPN.2C CG DNA models, respectively. Additionally, specific energy values for two representative conformations are listed in Tables 3 and 4 for reference. Note that the original CafeMol implementation follows the asymmetric definition for cross-stacking potential proposed in the original 3SPN.2 model^20^. To ensure a consistent comparison, we symmetrized the CafeMol results by averaging the energies of two configurations: (i) the original conformation, and (ii) a strand-swapped variant representing an exchange of sense and antisense strands. The resulting mean values enable a direct comparison with symmetric potentials in OpenCafeMol and Mjolnir.

**Figure 4.**
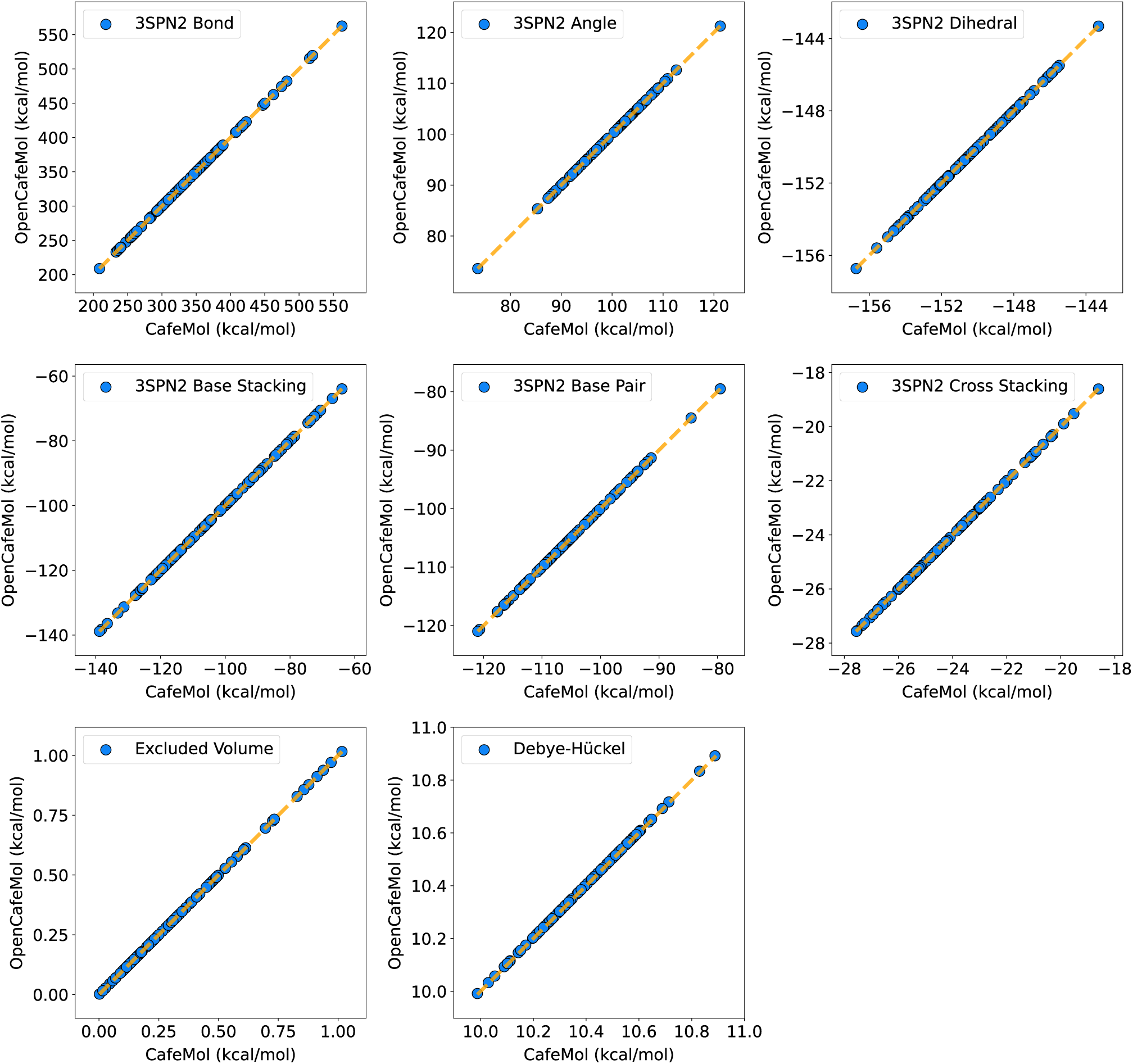
Comparison of individual energy terms of the 3SPN2 DNA model between CafeMol and OpenCafeMol for various DNA conformations.

**Figure 5.**
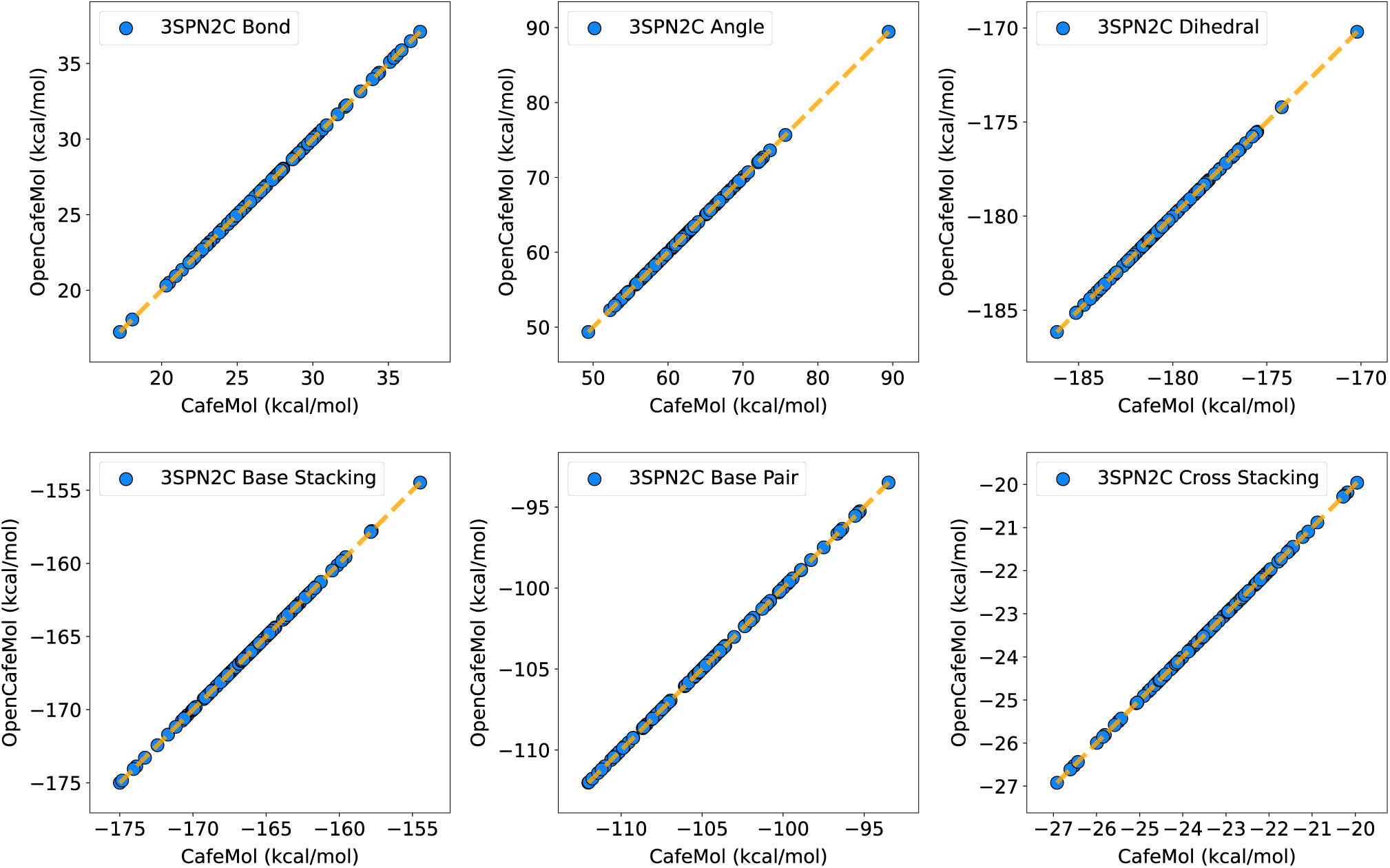
Comparison of individual energy terms of the 3SPN2C DNA model between CafeMol and OpenCafeMol for various DNA conformations.

**Table 3.** Comparison of energy values of the 3SPN2 DNA Model calculated from OpenCafeMol and Mjolnir.

| Potential<br>(kcal/mol) | OpenCafeMol<br>(conf. 1) | CafeMol<br>(conf. 1) | Mjolnir<br>(conf. 1) | OpenCafeMol<br>(conf. 2) | CafeMol<br>(conf. 2) | Mjolnir<br>(conf. 2) |
| --- | --- | --- | --- | --- | --- | --- |
| $V_{\text{Bond}}$ | 0.002016 | 0.002016 | 0.002016 | 25.18 | 25.18 | 25.18 |
| $V_{\text{Angle}}$ | 0.000315 | 0.000315 | 0.000315 | 55.03 | 55.03 | 55.03 |
| $V_{\text{Dihedral}}$ | -172.08 | -172.08 | -172.08 | -159.93 | -159.94 | -159.94 |
| $V_{\text{BaseStacking}}$ | -196.16 | -196.15 | -196.16 | -170.40 | -170.39 | -170.40 |
| $V_{\text{BasePair}}$<br>(localized) | -145.07<br>(-145.07) | -145.07 | -145.07 | -131.16<br>(-131.16) | -131.17 | -131.16 |
| $V_{\text{CrossStacking}}$<br>(localized) | -38.33<br>(-38.33) | -38.32 | -38.33 | -25.69<br>(-25.69) | -26.68 | -25.69 |
| $V_{\text{3SPN2 Excluded}}$ | 0.00 | 0.00 | 0.00 | 0.2726 | 0.2726 | 0.2726 |
| $V_{\text{Electrostatic}}$ | 11.22 | 11.22 | 11.22 | 10.44 | 10.44 | 10.44 |

**Table 4.** Comparison of energy values of the 3SPN2C DNA Model calculated from OpenCafeMol and Mjolnir.

| Potential<br>(kcal/mol) | OpenCafeMol<br>(conf. 1) | CafeMol<br>(conf.1) | Mjolnir<br>(conf. 1) | OpenCafeMol<br>(conf. 2) | CafeMol<br>(conf. 2) | Mjolnir<br>(conf. 2) |
| --- | --- | --- | --- | --- | --- | --- |
| $V_{\text{Bond}}$ | 165.88 | 165.88 | 165.88 | 215.29 | 215.29 | 215.29 |
| $V_{\text{Angle}}$ | 92.95 | 92.95 | 92.95 | 155.49 | 155.49 | 155.49 |
| $V_{\text{Dihedral}}$ | -192.54 | -192.54 | -192.54 | -183.33 | -183.32 | -183.33 |
| $V_{\text{BaseStacking}}$ | -160.97 | -160.97 | -160.97 | -137.61 | -137.60 | -137.61 |
| $V_{\text{BasePair}}$<br>(localized) | -122.05<br>(-122.05) | -122.05 | -122.05 | -101.63 | -101.63 | -101.63 |
| $V_{\text{CrossStacking}}$<br>(localized) | -25.67 | -25.66 | -25.67 | -15.34<br>(-15.34) | -15.34 | -15.34 |
| $V_{\text{3SPN2 Excluded}}$ | 0.00 | 0.00 | 0.00 | 0.2726 | 0.2726 | 0.2726 |
| $V_{\text{Electrostatic}}$ | 11.22 | 11.22 | 11.22 | 10.44 | 10.44 | 10.44 |

The results demonstrate excellent agreement between the potential energies calculated by OpenCafeMol and the reference software. Furthermore, comparisons in Tables 3 and 4 confirm that the localized treatment of base-pair and cross-stacking interactions introduced in OpenCafeMol yields energy values virtually identical to the global calculation, validating the accuracy of this optimization for double-stranded DNA.

### Benchmark 1: DNA-only system

To evaluate the computational efficiency of our implementations, we performed benchmark simulations on systems consisting exclusively of DNA molecules. The initial configurations consist of DNA molecules arranged in a three-dimensional grid, each comprising 47 base pairs with an identical sequence (GGCGACGTGA TCACCAGATG ATGCTAGATG CTTTCCGAAG AGAGAGC). System sizes ranged from a single DNA molecule to a 5×5×5 (125 molecules), corresponding to approximately 280 to 35,000 CG particles. All benchmarks used the 3SPN.2C CG DNA model^21^. Each configuration was simulated for 1×10^5^ time steps across ten independent runs.

Figure 6 illustrates the performance scaling of GPU-based OpenCafeMol simulations compared to CPU-based CafeMol. The performance gain from GPU acceleration increases markedly with system size. For the largest system (35,000 particles), a single GPU run with OpenCafeMol achieved an approximately 200-fold speed-up over a single CPU core and a 40-fold improvement over an 8-core CPU execution (Fig. 6C). In contrast, for small systems (<1,000 particles), the GPU acceleration is negligible or even detrimental. This reduced efficiency in the small-system regime is attributed to insufficient GPU occupancy, where the computational workload per GPU thread is too small to fully exploit the device’s parallelism, making kernel launch overheads significant.

**Figure 6.**
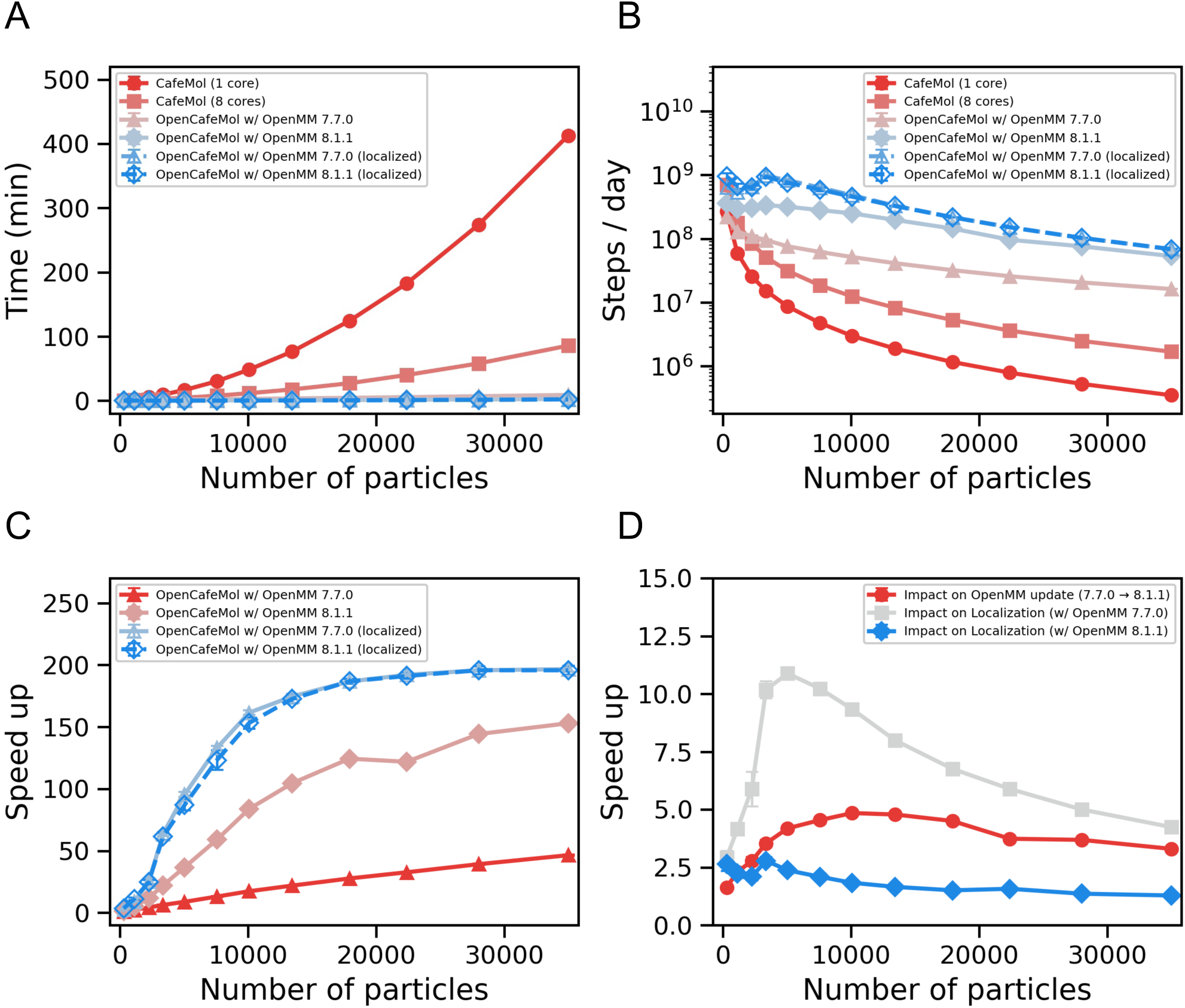
Performance benchmarking for DNA-only systems. (A) Wall-clock time required for 1×10^5^ steps. (B) Estimated throughput per day. (C) Speed-up compared to the single-core CafeMol calculations. (D) Impact of OpenMM version and interaction localization on performance. The red line indicates the speed-up from updating OpenMM 7.7.0 to 8.1.1. The gray and blue lines show the additional speed-up achieved by localizing base-pair and cross-stacking interactions within OpenMM 7.7.0 and 8.1.1, respectively.

We also investigated the impact of the underlying OpenMM version on performance specifically comparing versions OpenMM 7.7.0^39^ and OpenMM 8.1.1^33^. As shown in Fig. 6D, OpenMM 8.1.1 achieves up to a 5-fold performance improvement over OpenMM 7.7.0. This performance improvement is mainly due to an optimization of the CustomHBondForce kernel code used for base-pairing and cross-stacking interactions (see discussion on GitHub^34^).

Furthermore, we demonstrated that restricting base-pairing and cross-stacking calculations to only local Watson-Crick pairs and their neighbors significantly enhances performance (Fig. 6D). This localized interaction scheme yielded speed-ups of up to 10-fold with OpenMM 7.7.0. Even with the optimized kernels in OpenMM 8.1.1, the localization strategy provided an additional 2-fold acceleration, highlighting its distinct contribution to efficiency.

### Benchmark 2: nucleosome as an example of protein-DNA complexes

To evaluate the performance gains of our GPU-accelerated implementation in systems containing both proteins and DNA, we selected nucleosomes as a test case. The initial structures were derived from the 1KX5 crystal structure^43^, which consists of a histone octamer and 147 base pairs of DNA. In the simulations, AICG2+ force field^16^ was used for the histone octamer and the 3SPN.2C model^21^ was used for DNA. Debye-Hückel electrostatics, excluded volume, and hydrogen bond type interactions^23^ were considered for protein-DNA interactions (see our previous work for more details^25^.). The ionic concentration of the system was set to 150 mM. We investigated performance scaling by arranging nucleosomes in a three-dimensional grid to vary the system size. The systems ranged from a single nucleosome (1,860 particles where 880 particles for DNA and 980 particles for proteins) to 18 nucleosomes (3×3×2 grid; 33,480 particles in total) (Fig. 7A). Ten independent simulations of 1×10^5^ time steps were performed for each configuration.

**Figure 7.**
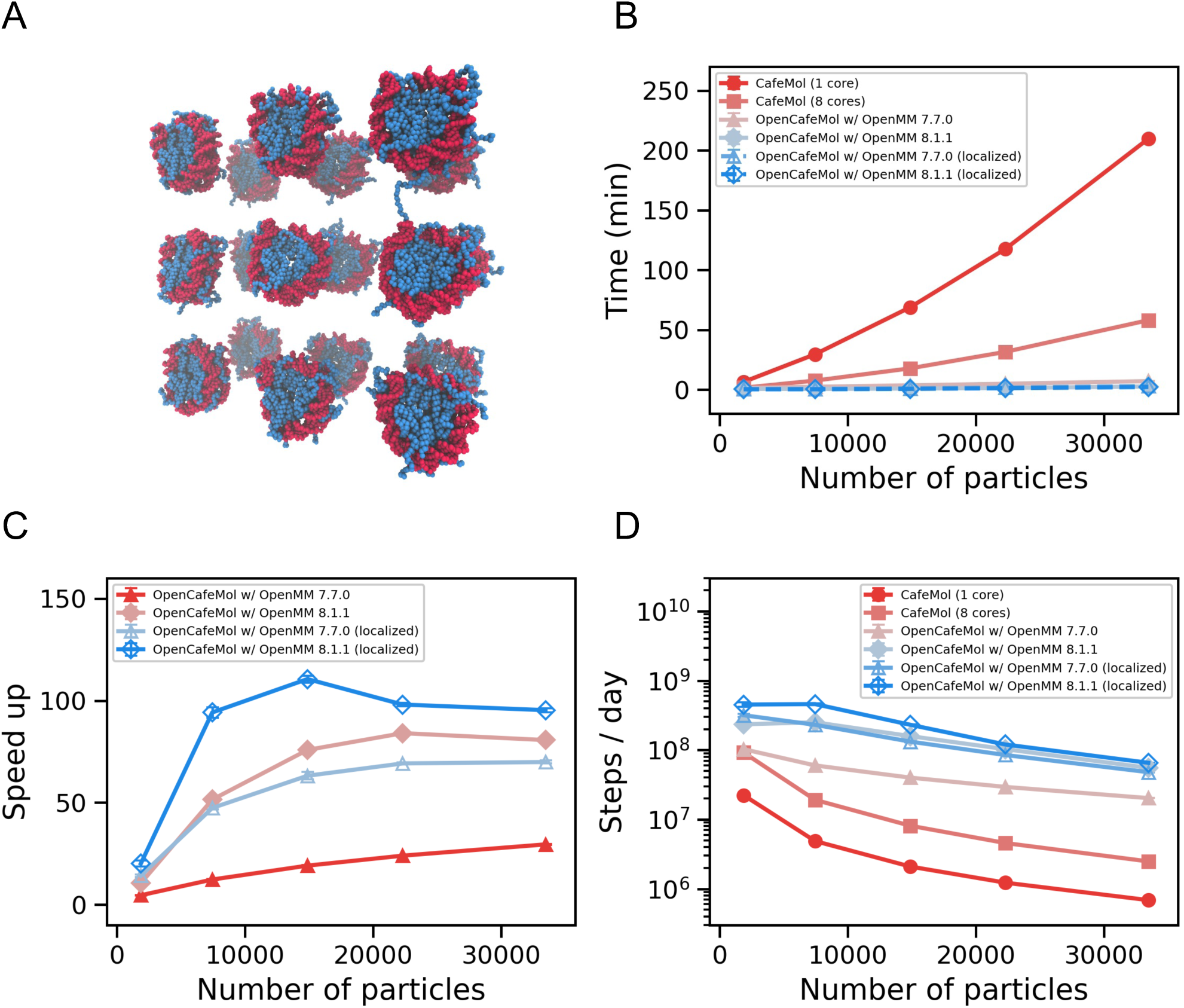
Performance benchmarking for nucleosome systems. (A) A structure of the coarse-grained nucleosomes in a 3×3×2 grid. (B) Wall-clock time required for 1×10^5^ steps. (C) Speed-up compared to a single-core CafeMol baseline. (D) Estimated simulation throughput per day.

Figure 7B shows the runtime for a time integration of 1×10^5^ steps for various numbers of nucleosomes. Consistent with the DNA-only systems (Figs. 6A-C), the performance gain from the GPU acceleration increased markedly with system size. Notably, the GPU-based simulations achieve up to 100-fold speedup compared to the CPU-based simulations (Fig. 7C). Figure 7D shows the estimated simulation throughput per day. Previous studies of nucleosomes using CPU-based CafeMol often required time integrations of up to 1×10^8^ steps^25–27^. Simulating a widely studied 12-nucleosome chromatin array^44–46^ (about 24,000 particles in the CG representation) at this timescale takes about 22 days using CafeMol on an 8-core CPU. In contrast, OpenCafeMol with GPU acceleration can complete the same task in less than a day. This significant efficiency gain enables the routine exploration of long-timescale biological phenomena that were previously computationally prohibitive.

### Benchmark 3: Smc-ScpA complex with 800bp double-stranded DNA

As an additional benchmark for DNA-protein complexes, we selected a system consisting of an archaeal SMC complex and an 800 bp double-stranded DNA (Fig. 8 inset). This system comprises 7,350 CG particles (4,798 particles for DNA and 2,552 particles for proteins). The SMC complex functions as a molecular motor, converting large-scale ATP-dependent conformational changes into mechanical work for DNA loop extrusion or DNA translocation to organize genome architecture^47–49^. A notable feature of the SMC complex is its slow ATP hydrolysis rate, occurring on the millisecond timescale^50^, as well as its large-scale conformational and DNA structure change. Observing the associated DNA dynamics requires longer-timescale simulations.

**Figure 8.**
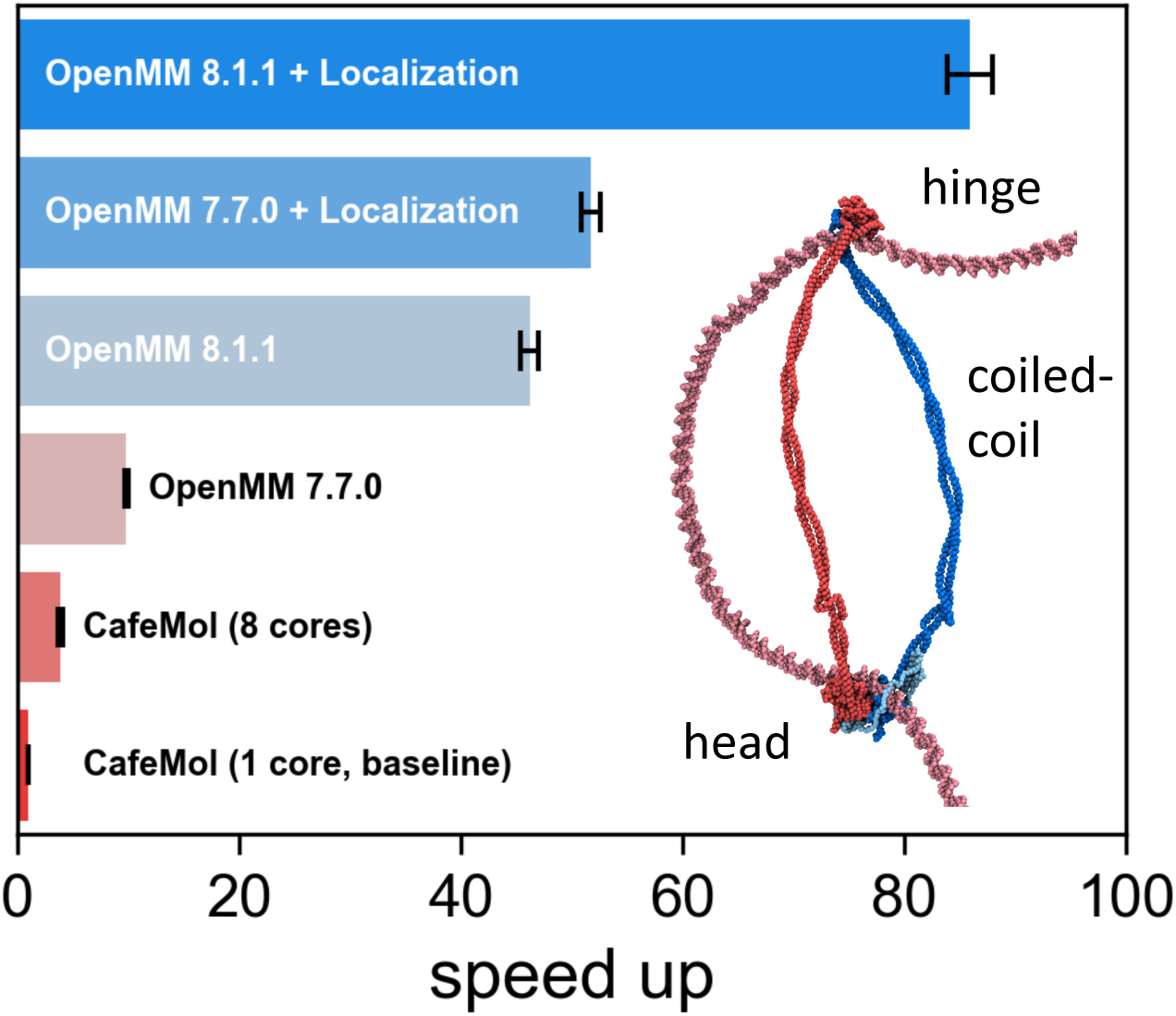
Benchmarking of Smc-ScpA complex with segment-captured 800-bp double-strand DNA. (inset) Structure of an archaeal Smc-ScpA complex. Smc homo dimer in red and blue; ScpA (kleisin) in cyan; DNA in pink.

Figure 8 illustrates the speed-up achieved by OpenCafeMol with GPU acceleration compared to the runtime of CafeMol on a single CPU core as the baseline. When base-pair and cross-stacking interactions were localized, OpenCafeMol simulations with GPU acceleration achieved an ∼84-fold speed-up. Even without such localization, a ∼47-fold speed-up was still achieved. In our recent CG simulation study^31^, reproducing DNA translocation through a complete ATP hydrolysis cycle of the SMC complex required time integration over 5×10^8^ steps, which took approximately 90 days on a single CPU core or 25 days on eight CPU cores using CafeMol. The performance improvement from the GPU implementation reduces the simulation time from over months or weeks on a CPU to just 1–2 days, facilitating efficient exploration of long-timescale dynamics in large chromatin-associated systems.

### Obstacle bypass by Smc-ScpA complex

To demonstrate the capability of OpenCafeMol to handle biologically relevant events involving large-scale structural changes, we simulated a prokaryotic Smc-ScpA complex encountering a protein obstacle on DNA. The prokaryotic Smc consists of an ATPase head domain, a hinge domain, and coiled-coil connecting the head and hinge domains, and forms a homo-dimer with its hinge domain. Two head domains are connected by ScpA, which is a member of kleisin family. Many SMC complexes function as ATP-driven molecular motors that extrude DNA loop or translocate along DNA. This activity is robust against physical barriers; for examples, i*n vivo* studies have shown that SMC translocation rates remain unaffected even in the presence of excess nucleoid-associated proteins^51^, and single-molecule experiments have revealed that SMC complexes can bypass large obstacles ranging from nucleosomes to nanoparticles^52^. To investigate the mechanistic basis of this bypass, we constructed a system comprising the *Pyrococcus yayanosii* Smc-ScpA complex, an 800-bp double-stranded DNA, and the Leucine-responsive regulatory protein (LrpA) as a DNA-bound obstacle. The CG parameters for Smc-ScpA and protein-DNA interactions were adopted from our previous study^31^. The CG parameter for LrpA is derived from a homology model based on *Pyrococcus furiosus* LrpA crystal structure (PDB: 1I1G)^53^. The structure of the LrpA-DNA complex was predicted using AlphaFold3^54^, and the specific protein-DNA interactions were modeled using Gō-like contact potentials based on the predicted complex structure.

We analyzed the time-evolution of protein-DNA contact to elucidate the dynamic process of obstacle bypass within the framework of the segment-capture mechanism (Fig. 9A and Movie1)^31,55–57^. The simulation mimics the ATP hydrolysis cycle by switching the potential energy function. In the initial phase (0 to 1×10^8^ steps), the SMC complex was maintained in the disengaged (apo) state. DNA contacts were primarily localized to kleisin subunit, confirming that the DNA was topologically entrapped within the kleisin ring. At 1×10^8^ steps (indicated by the dashed vertical line), we switched the potential energy functions to the engaged (ATP-bound) state to trigger the conformational transition. Following this transition, the ATPase heads rapidly engaged each other. Subsequently, we observed the progressive expansion of DNA contacts along the coiled-coil arms (Fig. 9A, green dots). This expansion indicates the successful initiation and growth of a DNA loop via the segment capture process. The contact points extended from approximately 400 bp to nearly 700 bp over the next 3×10^8^ steps, demonstrating continuous DNA loop growth. Importantly, LrpA-bound DNA segment did not impede loop formation but was successfully inserted into the internal compartment of the SMC ring formed by the coiled-coils. This observation suggests that the SMC complex can accommodate and bypass obstacles of this size via the segment-capture mechanism.

**Figure 9.**
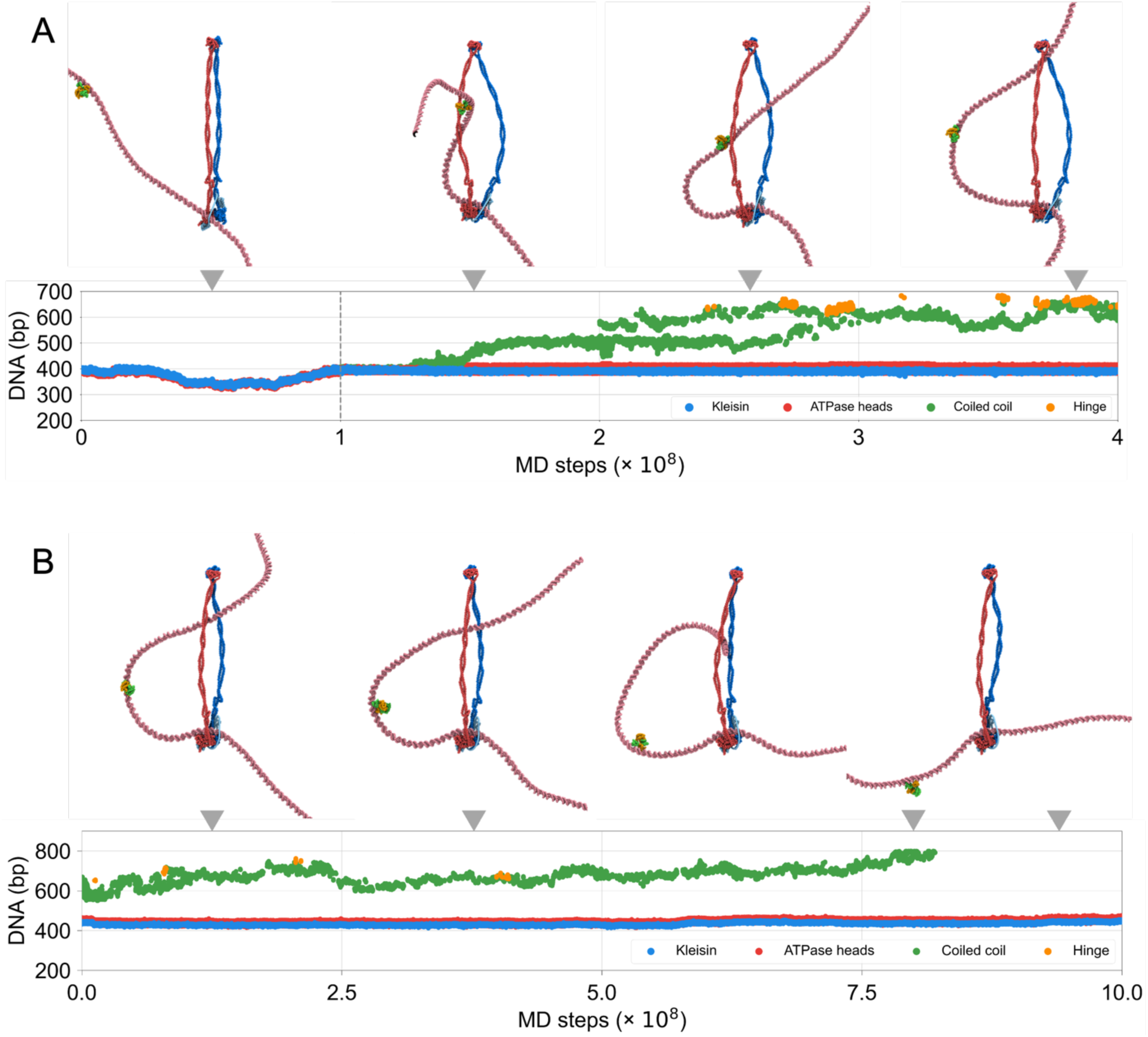
DNA segment capture simulations for SMC-ScpA complex. (A) Dynamics of obstacle bypass via DNA segment capture mechanism. (B) Expansion of the captured DNA loop length within the SMC ring in the engaged state. The data points represent the contact positions of each structural unit along the DNA in base pairs.

To explore the long-term dynamics captured DNA loop growth, we extended the simulation in the engaged state to 1×10^9^ steps. Fig. 9B and Movie2 show a trajectory revealing further expansion of the captured DNA segment. The DNA contact points along the coiled-coil arms migrated from approximately 600 bp to 800 bp over the course of the simulation. This indicates that the DNA loop within the SMC ring continued to grow via thermal fluctuations while being stabilized by the engaged ATPase heads. The loop size gradually increased to the physical length of the DNA substrate, and eventually, the distal end of the DNA passed through the hinge region, resulting in escape from the SMC complex. These results suggest that the DNA loop can continue to grow via thermal fluctuations as long as the complex remains in the engaged state, providing a potential explanation for the large step size exceeding the average of ∼200 bp reported in previous experimental measurements^58–60^.

## CONCLUSIONS

In this study, we successfully extended OpenCafeMol^15^, a GPU-accelerated residue-level CG MD simulator, to support 3SPN.2 and 3SPN.2C CG DNA models^20,21,23^ and hydrogen-bond-type potential^23^, enabling efficient simulation of DNA-protein interactions on GPU platforms. Our implementation achieved significant performance improvements, with up to 200-fold speedups for DNA-only systems (Fig. 6) and 100-fold speedups for nucleosome and large protein-DNA complexes compared to conventional CPU-based simulations (Figs. 7 and 8). These performance improvements were made possible through the optimization of many-body interactions, particularly base-pairing and cross-stacking potentials.

To further enhance computational efficiency, we introduced localization restriction of base-pairing and cross-stacking interactions to local base pairs and their neighbors. While this approach significantly reduces computational cost and enables efficient simulations of double-strand DNA, it inherently limits the generality of the 3SPN.2 and 3SPN.2C models. Specifically, this restriction hinders the applicability of these models for processes like hybridization or other scenarios involving single-stranded DNA or sliding of single-strand with respect to the other strand. Future developments could address this limitation by implementing more versatile algorithms, such as those based on neighbor lists or similar spatial partitioning algorithms. These methods have the potential to retain the general applicability of 3SPN.2 and 3SPN.2C models while achieving comparable or even greater computational efficiency.

In summary, this work provides a powerful computational tool for chromatin biology studies, addressing the increasing demand for high-performance simulations of DNA-containing biomolecular systems. We believe that the advancements presented here will be useful for deeper exploration of a wide range of chromatin-related processes.

## ACKNOWLEDGEMENT

We would like to thank Fritz Nagae for reporting bugs and technical support. We also would like to thank Giovanni B. Brandani for providing CafeMol input for the nucleosome system. We used the supercomputer at the Academic Center for Computing and Media Studies (Kyoto University). This study was supported by JSPS KAKENHI, Grant Number 20H05934 (S.T.), 21H02441 (S.T.), 24K01991 (S.T.), and by the MEXT grants JPMXP1020230119 as “Program for Promoting Researches on the Supercomputer Fugaku” (S.T.).

## CODE AVAILABILITY

The source code are publicly available at https://github.com/yutakasi634/OpenCafeMol.

## AUTHOR CONTRIBUTION

M.Y., Y.M., and S.T. conceived the project; M.Y., Y.M., and T.N. developed the code. M.Y. performed the simulations, analyzed the data, and assembled the figures; M.Y. and S.T. discussed the results and wrote the manuscript.

## REFERENCES

(1) Saunders, M. G.; Voth, G. A. Coarse-Graining Methods for Computational Biology. Annu. Rev. Biophys. 2013, 42 (1), 73‒93. 10.1146/annurev-biophys-083012-130348.

(2) Noid, W. G. Perspective: Coarse-Grained Models for Biomolecular Systems. J. Chem. Phys. 2013, 139 (9), 090901. 10.1063/1.4818908.

(3) Kmiecik, S.; Gront, D.; Kolinski, M.; Wieteska, L.; Dawid, A. E.; Kolinski, A. Coarse-Grained Protein Models and Their Applications. Chem. Rev. 2016, 116 (14), 7898‒7936. 10.1021/acs.chemrev.6b00163.

(4) Takada, S.; Kanada, R.; Tan, C.; Terakawa, T.; Li, W.; Kenzaki, H. Modeling Structural Dynamics of Biomolecular Complexes by Coarse-Grained Molecular Simulations. Acc. Chem. Res. 2015, 48 (12), 3026‒3035. 10.1021/acs.accounts.5b00338.

(5) Jung, J.; Mori, T.; Kobayashi, C.; Matsunaga, Y.; Yoda, T.; Feig, M.; Sugita, Y. GENESIS: A Hybrid-Parallel and Multi-Scale Molecular Dynamics Simulator with Enhanced Sampling Algorithms for Biomolecular and Cellular Simulations. Wiley Interdiscip. Rev. Comput. Mol. Sci. 2015, 5 (4), 310‒323. 10.1002/wcms.1220.

(6) Kobayashi, C.; Jung, J.; Matsunaga, Y.; Mori, T.; Ando, T.; Tamura, K.; Kamiya, M.; Sugita, Y. GENESIS 1.1: A Hybrid-Parallel Molecular Dynamics Simulator with Enhanced Sampling Algorithms on Multiple Computational Platforms. J. Comput. Chem. 2017, 38 (25), 2193‒2206. 10.1002/jcc.24874.

(7) Jung, J.; Yagi, K.; Tan, C.; Oshima, H.; Mori, T.; Yu, I.; Matsunaga, Y.; Kobayashi, C.; Ito, S.; Ugarte La Torre, D.; Sugita, Y. GENESIS 2.1: High-Performance Molecular Dynamics Software for Enhanced Sampling and Free-Energy Calculations for Atomistic, Coarse-Grained, and Quantum Mechanics/Molecular Mechanics Models. J. Phys. Chem. B 2024. 10.1021/acs.jpcb.4c02096.

(8) Abraham, M. J.; Murtola, T.; Schulz, R.; Páll, S.; Smith, J. C.; Hess, B.; Lindahl, E. GROMACS: High Performance Molecular Simulations through Multi-Level Parallelism from Laptops to Supercomputers. SoftwareX 2015, 1-2, 19‒25. 10.1016/j.softx.2015.06.001.

(9) Götz, A. W.; Williamson, M. J.; Xu, D.; Poole, D.; Le Grand, S.; Walker, R. C. Routine Microsecond Molecular Dynamics Simulations with AMBER on GPUs. 1. Generalized Born. J. Chem. Theory Comput. 2012, 8 (5), 1542‒1555. 10.1021/ct200909j.

(10) Salomon-Ferrer, R.; Götz, A. W.; Poole, D.; Le Grand, S.; Walker, R. C. Routine Microsecond Molecular Dynamics Simulations with AMBER on GPUs. 2. Explicit Solvent Particle Mesh Ewald. J. Chem. Theory Comput. 2013, 9 (9), 3878‒3888. 10.1021/ct400314y.

(11) Le Grand, S.; Götz, A. W.; Walker, R. C. SPFP: Speed without Compromise - A Mixed Precision Model for GPU Accelerated Molecular Dynamics Simulations. Comput. Phys. Commun. 2013, 184 (2), 374‒380. 10.1016/j.cpc.2012.09.022.

(12) Eastman, P.; Pande, V. OpenMM: A Hardware-Independent Framework for Molecular Simulations. Comput. Sci. Eng. 2010, 12 (4), 34‒39. 10.1109/MCSE.2010.27.

(13) Phillips, J. C.; Hardy, D. J.; Maia, J. D. C.; Stone, J. E.; Ribeiro, J. V.; Bernardi, R. C.; Buch, R.; Fiorin, G.; Hénin, J.; Jiang, W.; McGreevy, R.; Melo, M. C. R.; Radak, B. K.; Skeel, R. D.; Singharoy, A.; Wang, Y.; Roux, B.; Aksimentiev, A.; Luthey-Schulten, Z.; Kalé, L. V.; Schulten, K.; Chipot, C.; Tajkhorshid, E. Scalable Molecular Dynamics on CPU and GPU Architectures with NAMD. Journal of Chemical Physics 2020, 153 (4). 10.1063/5.0014475.

(14) Thompson, A. P.; Aktulga, H. M.; Berger, R.; Bolintineanu, D. S.; Brown, W. M.; Crozier, P. S.; in ’t Veld, P. J.; Kohlmeyer, A.; Moore, S. G.; Nguyen, T. D.; Shan, R.; Stevens, M. J.; Tranchida, J.; Trott, C.; Plimpton, S. J. LAMMPS - a Flexible Simulation Tool for Particle-Based Materials Modeling at the Atomic, Meso, and Continuum Scales. Comput. Phys. Commun. 2022, 271. 10.1016/j.cpc.2021.108171.

(15) Murata, Y.; Niina, T.; Takada, S. OpenCafeMol: A Coarse-Grained Biomolecular Simulator on GPU with Its Application to Vesicle Fusion. Biophys. J. 2025. 10.1016/j.bpj.2025.07.012.

(16) Li, W.; Wang, W.; Takada, S. Energy Landscape Views for Interplays among Folding, Binding, and Allostery of Calmodulin Domains. Proceedings of the National Academy of Sciences 2014, 111 (29), 10550‒10555. 10.1073/pnas.1402768111.

(17) Dignon, G. L.; Zheng, W.; Kim, Y. C.; Best, R. B.; Mittal, J. Sequence Determinants of Protein Phase Behavior from a Coarse-Grained Model. PLoS Comput. Biol. 2018, 14 (1), 1‒23. 10.1371/journal.pcbi.1005941.

(18) Tesei, G.; Schulze, T. K.; Crehuet, R.; Lindorff-Larsen, K. Accurate Model of Liquid‒Liquid Phase Behavior of Intrinsically Disordered Proteins from Optimization of Single-Chain Properties. Proceedings of the National Academy of Sciences 2021, 118 (44), 1‒9. 10.1073/pnas.2111696118.

(19) La Torre, D. U.; Takada, S. Coarse-Grained Implicit Solvent Lipid Force Field with a Compatible Resolution to the C α Protein Representation. Journal of Chemical Physics 2020, 153 (20). 10.1063/5.0026342.

(20) Hinckley, D. M.; Freeman, G. S.; Whitmer, J. K.; de Pablo, J. J. An Experimentally-Informed Coarse-Grained 3-Site-per-Nucleotide Model of DNA: Structure, Thermodynamics, and Dynamics of Hybridization. J. Chem. Phys. 2013, 139 (14), 144903. 10.1063/1.4822042.

(21) Freeman, G. S.; Hinckley, D. M.; Lequieu, J. P.; Whitmer, J. K.; de Pablo, J. J. Coarse-Grained Modeling of DNA Curvature. J. Chem. Phys. 2014, 141 (16), 165103. 10.1063/1.4897649.

(22) Knotts, T. A.; Rathore, N.; Schwartz, D. C.; de Pablo, J. J. A Coarse Grain Model for DNA. J. Chem. Phys. 2007, 126 (8), 084901. 10.1063/1.2431804.

(23) Niina, T.; Brandani, G. B.; Tan, C.; Takada, S. Sequence-Dependent Nucleosome Sliding in Rotation-Coupled and Uncoupled Modes Revealed by Molecular Simulations. PLoS Comput. Biol. 2017, 13 (12), e1005880. 10.1371/journal.pcbi.1005880.

(24) Brandani, G. B.; Gopi, S.; Yamauchi, M.; Takada, S. Molecular Dynamics Simulations for the Study of Chromatin Biology. Curr. Opin. Struct. Biol. 2022, 77 (Md), 102485. 10.1016/j.sbi.2022.102485.

(25) Brandani, G. B.; Niina, T.; Tan, C.; Takada, S. DNA Sliding in Nucleosomes via Twist Defect Propagation Revealed by Molecular Simulations. Nucleic Acids Res. 2018, 46 (6), 2788‒2801. 10.1093/nar/gky158.

(26) Brandani, G. B.; Takada, S. Chromatin Remodelers Couple Inchworm Motion with Twist-Defect Formation to Slide Nucleosomal DNA. PLoS Comput. Biol. 2018, 14 (11), e1006512. 10.1371/journal.pcbi.1006512.

(27) Tan, C.; Takada, S. Nucleosome Allostery in Pioneer Transcription Factor Binding. Proc. Natl. Acad. Sci. U. S. A. 2020, 117 (34), 20586‒20596. 10.1073/pnas.2005500117.

(28) Shino, G.; Takada, S. Modeling DNA Opening in the Eukaryotic Transcription Initiation Complexes via Coarse-Grained Models. Front. Mol. Biosci. 2021, 8. 10.3389/fmolb.2021.772486.

(29) Nagae, F.; Murata, Y.; Yamauchi, M.; Takada, S.; Terakawa, T. Mechanistic Models of Asymmetric Hand-over-Hand Translocation and Nucleosome Navigation by CMG Helicase. Nat. Commun. 2025, 16 (1), 10304. 10.1038/s41467-025-65232-x.

(30) Nagae, F.; Murayama, Y.; Terakawa, T. Molecular Mechanism of Parental H3/H4 Recycling at a Replication Fork. Nat. Commun. 2024, 15 (1), 9485. 10.1038/s41467-024-53187-4.

(31) Yamauchi, M.; Brandani, G. B.; Terakawa, T.; Takada, S. SMC Complex Unidirectionally Translocates DNA by Coupling Segment Capture with an Asymmetric Kleisin Path. eLife. July 16, 2025. 10.7554/eLife.106752.1.

(32) Lu, W.; Bueno, C.; Schafer, N. P.; Moller, J.; Jin, S.; Chen, X.; Chen, M.; Gu, X.; Davtyan, A.; de Pablo, J. J.; Wolynes, P. G. OpenAWSEM with Open3SPN2: A Fast, Flexible, and Accessible Framework for Large-Scale Coarse-Grained Biomolecular Simulations. PLoS Comput. Biol. 2021, 17 (2), e1008308. 10.1371/journal.pcbi.1008308.

(33) Eastman, P.; Galvelis, R.; Peláez, R. P.; Abreu, C. R. A.; Farr, S. E.; Gallicchio, E.; Gorenko, A.; Henry, M. M.; Hu, F.; Huang, J.; Krämer, A.; Michel, J.; Mitchell, J. A.; Pande, V. S.; Rodrigues, J. P.; Rodriguez-Guerra, J.; Simmonett, A. C.; Singh, S.; Swails, J.; Turner, P.; Wang, Y.; Zhang, I.; Chodera, J. D.; De Fabritiis, G.; Markland, T. E. OpenMM 8: Molecular Dynamics Simulation with Machine Learning Potentials. J. Phys. Chem. B 2024, 128 (1), 109‒116. 10.1021/acs.jpcb.3c06662.

(34) Peter Eastman. Faster implementation of CustomHbondForce. GitHub. https://github.com/openmm/openmm/pull/4060.

(35) Lu, X.-J.; Wilma K. Olson. 3DNA: A Software Package for the Analysis, Rebuilding and Visualization of Three-Dimensional Nucleic Acid Structures. Nucleic Acids Res. 2003, 31 (17), 5108‒5121. 10.1093/nar/gkg680.

(36) Lu, X.-J.; Olson, W. K. 3DNA: A Versatile, Integrated Software System for the Analysis, Rebuilding and Visualization of Three-Dimensional Nucleic-Acid Structures. Nat. Protoc. 2008, 3 (7), 1213‒1227. 10.1038/nprot.2008.104.

(37) Toru Niina. Mjolnir: General Purpose Coarse-Grained Molecular Dynamics Simulation Package. https://github.com/Mjolnir-MD/Mjolnir (accessed 2025-04-12).

(38) Tan, C.; Terakawa, T.; Takada, S. Dynamic Coupling among Protein Binding, Sliding, and DNA Bending Revealed by Molecular Dynamics. J. Am. Chem. Soc. 2016, 138 (27), 8512‒8522. 10.1021/jacs.6b03729.

(39) Eastman, P.; Swails, J.; Chodera, J. D.; McGibbon, R. T.; Zhao, Y.; Beauchamp, K. A.; Wang, L.-P.; Simmonett, A. C.; Harrigan, M. P.; Stern, C. D.; Wiewiora, R. P.; Brooks, B. R.; Pande, V. S. OpenMM 7: Rapid Development of High Performance Algorithms for Molecular Dynamics. PLoS Comput. Biol. 2017, 13 (7), e1005659. 10.1371/journal.pcbi.1005659.

(40) Eastman, P.; Pande, V. Accelerating Development and Execution Speed with Just-in-Time GPU Code Generation. In GPU Computing Gems Jade Edition; Elsevier, 2012; pp 399‒407. 10.1016/B978-0-12-385963-1.00029-0.

(41) Kenzaki, H.; Koga, N.; Hori, N.; Kanada, R.; Li, W.; Okazaki, K.; Yao, X.-Q.; Takada, S. CafeMol: A Coarse-Grained Biomolecular Simulator for Simulating Proteins at Work. J. Chem. Theory Comput. 2011, 7 (6), 1979‒1989. 10.1021/ct2001045.

(42) Honeycutt, J. D.; Thirumalai, D. The Nature of Folded States of Globular Proteins. Biopolymers 1992, 32 (6), 695‒709. 10.1002/bip.360320610.

(43) Davey, C. A.; Sargent, D. F.; Luger, K.; Maeder, A. W.; Richmond, T. J. Solvent Mediated Interactions in the Structure of the Nucleosome Core Particle at 1.9 Å Resolution. J. Mol. Biol. 2002, 319 (5), 1097‒1113. 10.1016/S0022-2836(02)00386-8.

(44) Correll, S. J.; Schubert, M. H.; Grigoryev, S. A. Short Nucleosome Repeats Impose Rotational Modulations on Chromatin Fibre Folding. EMBO Journal 2012, 31 (10), 2416‒2426. 10.1038/emboj.2012.80.

(45) Song, F.; Chen, P.; Sun, D.; Wang, M.; Dong, L.; Liang, D.; Xu, R.-M.; Zhu, P.; Li, G. Cryo-EM Study of the Chromatin Fiber Reveals a Double Helix Twisted by Tetranucleosomal Units. Science (1979). 2014, 344 (6182), 376‒380. 10.1126/science.1251413.

(46) Farr, S. E.; Woods, E. J.; Joseph, J. A.; Garaizar, A.; Collepardo-Guevara, R. Nucleosome Plasticity Is a Critical Element of Chromatin Liquid‒Liquid Phase Separation and Multivalent Nucleosome Interactions. Nat. Commun. 2021, 12 (1), 1‒17. 10.1038/s41467-021-23090-3.

(47) Yatskevich, S.; Rhodes, J.; Nasmyth, K. Organization of Chromosomal DNA by SMC Complexes. Annu. Rev. Genet. 2019, 53 (1), 445‒482. 10.1146/annurev-genet-112618-043633.

(48) Davidson, I. F.; Peters, J.-M. Genome Folding through Loop Extrusion by SMC Complexes. Nat. Rev. Mol. Cell Biol. 2021, 22 (7), 445‒464. 10.1038/s41580-021-00349-7.

(49) Kim, E.; Barth, R.; Dekker, C. Looping the Genome with SMC Complexes. Annu. Rev. Biochem. 2023, 92 (1), 15‒41. 10.1146/annurev-biochem-032620-110506.

(50) Hassler, M.; Shaltiel, I. A.; Haering, C. H. Towards a Unified Model of SMC Complex Function. Current Biology 2018, 28 (21), R1266‒R1281. 10.1016/j.cub.2018.08.034.

(51) Ren, Z.; Way, L. E.; Wang, X. SMC Translocation Is Unaffected by an Excess of Nucleoid Associated Proteins in Vivo. Sci. Rep. 2025, 15 (1). 10.1038/s41598-025-86946-4.

(52) Pradhan, B.; Barth, R.; Kim, E.; Davidson, I. F.; Bauer, B.; van Laar, T.; Yang, W.; Ryu, J.-K.; van der Torre, J.; Peters, J.-M.; Dekker, C. SMC Complexes Can Traverse Physical Roadblocks Bigger than Their Ring Size. Cell Rep. 2022, 41 (3), 111491. 10.1016/j.celrep.2022.111491.

(53) Leonard, P. M.; Smits, S. H. J.; Sedelnikova, S. E.; Brinkman, A. B.; De Vos, W. M.; Van Der Oost, J.; Rice, D. W.; Rafferty, J. B. Crystal Structure of the Lrp-like Transcriptional Regulator from the Archaeon Pyrococcus Furiosus. EMBO Journal 2001, 20 (5). 10.1093/emboj/20.5.990.

(54) Abramson, J.; Adler, J.; Dunger, J.; Evans, R.; Green, T.; Pritzel, A.; Ronneberger, O.; Willmore, L.; Ballard, A. J.; Bambrick, J.; Bodenstein, S. W.; Evans, D. A.; Hung, C. C.; O’Neill, M.; Reiman, D.; Tunyasuvunakool, K.; Wu, Z.; Žemgulytė, A.; Arvaniti, E.; Beattie, C.; Bertolli, O.; Bridgland, A.; Cherepanov, A.; Congreve, M.; Cowen-Rivers, A. I.; Cowie, A.; Figurnov, M.; Fuchs, F. B.; Gladman, H.; Jain, R.; Khan, Y. A.; Low, C. M. R.; Perlin, K.; Potapenko, A.; Savy, P.; Singh, S.; Stecula, A.; Thillaisundaram, A.; Tong, C.; Yakneen, S.; Zhong, E. D.; Zielinski, M.; Žídek, A.; Bapst, V.; Kohli, P.; Jaderberg, M.; Hassabis, D.; Jumper, J. M. Accurate Structure Prediction of Biomolecular Interactions with AlphaFold 3. Nature 2024, 630 (8016), 493‒500. 10.1038/s41586-024-07487-w.

(55) Diebold-Durand, M.-L.; Lee, H.; Ruiz Avila, L. B.; Noh, H.; Shin, H.-C.; Im, H.; Bock, F. P.; Bürmann, F.; Durand, A.; Basfeld, A.; Ham, S.; Basquin, J.; Oh, B.-H.; Gruber, S. Structure of Full-Length SMC and Rearrangements Required for Chromosome Organization. Mol. Cell 2017, 67 (2), 334–347.e5. 10.1016/j.molcel.2017.06.010.

(56) Marko, J. F.; De Los Rios, P.; Barducci, A.; Gruber, S. DNA-Segment-Capture Model for Loop Extrusion by Structural Maintenance of Chromosome (SMC) Protein Complexes. Nucleic Acids Res. 2019, 47 (13), 6956‒6972. 10.1093/nar/gkz497.

(57) Nomidis, S. K.; Carlon, E.; Gruber, S.; Marko, J. F. DNA Tension-Modulated Translocation and Loop Extrusion by SMC Complexes Revealed by Molecular Dynamics Simulations. Nucleic Acids Res. 2022, 50 (9), 4974‒4987. 10.1093/nar/gkac268.

(58) Wang, X.; Hughes, A. C.; Brandão, H. B.; Walker, B.; Lierz, C.; Cochran, J. C.; Oakley, M. G.; Kruse, A. C.; Rudner, D. Z. In Vivo Evidence for ATPase-Dependent DNA Translocation by the Bacillus Subtilis SMC Condensin Complex. Mol. Cell 2018, 71 (5), 841–847.e5. 10.1016/j.molcel.2018.07.006.

(59) Wang, X.; Brandão, H. B.; Le, T. B. K.; Laub, M. T.; Rudner, D. Z. Bacillus Subtilis SMC Complexes Juxtapose Chromosome Arms as They Travel from Origin to Terminus. Science (1979). 2017, 355 (6324), 524‒527. 10.1126/science.aai8982.

(60) Tran, N. T.; Laub, M. T.; Le, T. B. K. SMC Progressively Aligns Chromosomal Arms in Caulobacter Crescentus but Is Antagonized by Convergent Transcription. Cell Rep. 2017, 20 (9), 2057‒2071. 10.1016/j.celrep.2017.08.026.

